# A Hybrid Communication Pattern in Human Brain Structural Network Revealed by Evolutionary Computation

**DOI:** 10.1101/2021.11.25.469862

**Authors:** Quanmin Liang, Junji Ma, Xitian Chen, Zhengjia Dai, Ying Lin

## Abstract

The human brain functional connectivity network (FCN) is constrained and shaped by the communication processes in the structural connectivity network (SCN). The underlying communication mechanism thus becomes a critical issue for understanding the formation and organization of the FCN. A number of communication models supported by different routing strategies have been proposed, with shortest path (SP), random diffusion (DIF), and spatial navigation (NAV) as the most typical, respectively requiring network global knowledge, local knowledge, and both for path seeking. Yet these models all assumed every brain region to use one routing strategy uniformly, ignoring convergent evidence that supports the regional heterogeneity in both terms of biological substrates and functional roles. In this regard, the current study developed a hybrid communication model that allowed each brain region to choose a routing strategy from SP, DIF, and NAV independently. A genetic algorithm was designed to uncover the underlying region-wise hybrid routing strategy (namely HYB). The HYB was found to outperform the three typical routing strategies in predicting FCN and facilitating robust communication. Analyses on HYB further revealed that brain regions in lower-order functional modules inclined to route signals using global knowledge, while those in higher-order functional modules preferred DIF that requires only local knowledge. Compared to regions that used global knowledge for routing, regions using DIF had denser structural connections, participated in more functional modules, but played a less dominant role within modules. Together, our findings further evidenced that hybrid routing underpins efficient SCN communication and locally heterogeneous structure-function coupling.

## 1. Introduction

With the advance of magnetic resonance imaging (MRI) technology, the human neural system can be characterized as a complex network where gray matter regions (nodes) are interconnected by white-matter projections (links), namely the brain structural connectivity network (SCN) [1][2]. Previous studies have shown that the SCN possesses multiple non-random topological properties, including small-worldness [3], modularization [4], and rich-club organization [5], which facilitates communication among brain regions [6]. It has also been recognized that communication paths in the SCN (i.e., transmission routes of neural signals along white-matter pathways) are crucial for understanding how the functional connectivity network (FCN), where brain regions are connected in terms of temporal correlations, is constrained and shaped by the SCN [7][8]. A series of models concerning communication paths have been developed to predict FCN from SCN (e.g., shortest-path-based global efficiency and multi-path-based communicability) [9]-[11]. However, although the significance of SCN communication has been admitted and evidenced, the underlying strategy (i.e., how neural signals are routed through the white-matter pathways) remains an open question [12][13].

Initially, the exploration of the communication models in the SCN was mainly based on the shortest paths between brain regions [10][14][15], which is indeed fast and straightforward, effectively reducing the risk of signal loss [16]. However, the disadvantage is also obvious: every region must fully master the global network topology, which induces heavy informational costs [17]-[19]. In addition, the shortest-path routing utilizes only a small portion of links, resulting in a disproportionately large information flow over a small number of connections [20]. The network is thus prone to information congestion and targeted attack [21][22]. In contrast to the idea of optimal routing, some researchers proposed SCN communication strategies based on the concept of random walking [18]. Since random walkers rely purely on local topological information (e.g., connection strength) [19], the random-walking-based strategies remove the premise that all elements in the network need to know the global network topology and thus relieve the burden of informational cost. It can also reduce the information congestion phenomenon and improve the network robustness by balancing information flow on network connections. However, previous studies have shown that they may lead to longer transmission routes and thus slow down the transmission speed, reducing communication efficiency among brain regions [18].

Considering that neither optimal routing nor random walking fully support or explain the communication patterns in the human brain, the idea of combining local knowledge and spatial embedding was proposed, leading to a navigation routing strategy greedily guided by local adjacency and global spatial positions [6][23]. Recently, it was further suggested that the routing strategy in the human brain is probably on a spectrum spanning from pure utilization of global information (shortest-path routing) to pure utilization of local information (random walking) [19]. Notably, despite of combining different types of information, the above communication models still inherited a common premise: all brain regions use the same routing strategy. This premise, although significantly simplifies the communication dynamics in the brain network, ignores convergent evidence supporting that brain regions play different roles in the brain communication process. For example, the regions in the ‘rich-club’ organization, which are mutually and densely inter-connected, have been found more central in inter-region signal traffic, forming a backbone for global communication [24]. The resilience of human brain to neural lesions has been found related to the involvement of alternative paths, which were mostly supported by highly connected hub regions [25][26]. These differences in the communication roles of brain regions suggest that they may use different routing strategies to adapt their white-matter basis and serve respective purposes. However, very few studies have been explored the communication pattern using such routing strategy that brain regions may adopt different routing strategies

Based on previous findings, this study relaxed the premise of globally uniform routing and proposed a communication model supported by a region-wise hybrid routing framework that allowed brain regions to select routing strategies from three representatives: shortest path (SP), random diffusion (DIF), and spatial navigation (NAV). This hybrid framework, which can be viewed as a region-based discrete implementation for the idea of brain routing spectrum, allows brain regions to acquire only necessary knowledge for their own routing strategies (global topology for SP, local topology for DIF, global spatial embedding and local topology for NAV). Relying on the imaging data from a cohort of 88 healthy adults, a genetic algorithm [27], which is a model-free metaheuristic in the domain of evolutionary computation well-known for its global search capability, was designed to identify the region-wise routing pattern that best explained the variation in functional connectivity. Note that the proposed framework tolerates the extreme situation that all brain regions select the same routing strategy. By doing so, we allowed the genetic algorithm to fully explore potential routing hybridizations in comparison with their uniform counterparts. The advantages of the obtained routing hybridization were then examined from the angles of how well it supported the FCN and facilitated robust communication in the SCN. The regional choices on the routing strategies were also analyzed regarding their cognitive function and topological roles in brain networks.

## 2. Validation Analyses and Results

### Validation Analysis

The validation dataset (denoted as ‘BNU’) is a subset of the Connectivity-based Brain Imaging Research Database (C-BIRD) at Beijing Normal University and can be acquired from the publicly available data of Consortium for Reliability and Reproducibility (http://fcon_1000.projects.nitrc.org/indi/CoRR/html/bnu_1.html) [28]. The BNU dataset consisted of two MRI scan sessions from 57 healthy adults (male: 30; age: 19-30 years), completed over a time interval of approximately 6 weeks (40.94±4.51 days). In particular, the first scan session included two R-fMRI scans, T1, T2, and Diffusion-tensor images (DTI). Only the data of the first R-fMRI scan were used in the validation study. All participants were right-handed and had no history of neurological or psychiatric disorders. Written informed consent was obtained from each participant, and data collection was approved by the Institutional Review Board of the State Key Laboratory of Cognitive Neuroscience and Learning, Beijing Normal University. The BNU dataset was preprocessed using the same pipeline as in the main study. Two participants were excluded, one for excessive head motion (translation > 2 mm or rotation > 2°) and the other one for missing slices in R-fMRI data. The proposed GA was then applied to the group-average SCN of the 55 participants, and the resulting hybrid routing strategy (denoted as ‘B-HYB’) was analyzed to examine whether our main findings were reproducible in the following three aspects: advantage in reconstructing the FCN, support to robust communication, and the relationship between regional choices of routing strategies and their topological roles.

### Validation Results

Visual inspection on Fig. S2 found that the B-HYB strategy was similar with the HYB in terms of both spatial distribution and modular proportions. Further quantified analyses showed that between HYB and B-HYB, the spatial similarity was at a moderate level (Cohen’s Kappa coefficient = 0.383), whilst the modular proportions of the three classic routing strategies were all highly correlated (SP/DIF/NAV: *r* = 0.920/0.983/0.912, *p* < 0.01), suggesting that the hybrid routing pattern showed a convergent distribution at the modular level but was more diverged region-wise. The B-HYB also achieved a significantly higher correlation between structural routing efficiency and FC (*r* = 0.441, *p* < 0.001) than the three classic routing strategies did (SP/DIF/NAV: *r* = 0.366/0.338/0.376, *p* < 0.001; *t* = 9.639/14.517/10.054, *p* < 0.001; Fig. S3A), indicating that the advantage of region-wise hybrid routing in capturing the FCN was reproducible across datasets. The comparison between B-HYB and the three classic routing strategies on the distributions of normalized Bc and the results of targeted lesion simulation confirmed that region-wise hybrid routing consistently facilitated robust communication in the SCN (Fig. S3B-D). In addition, as in the main analyses, the B-HYB still retained over 70% of the optimal network global efficiency achieved by the SP strategy (gE = 0.097), suggesting that the ability of hybrid routing in striking a balance between robustness and efficiency was independent of datasets. Finally, the analyses on the topological features of B-HYB found consistent results with those of HYB. That is, the regions using DIF had significantly higher structural degree and functional Pc than those using SP and NAV, but significantly lower functional within-module Z-scores than those using SP (Fig. S4A-C). In addition, by classifying the regions using the same Pc-Z coordinate as in the main text, we found that the preference of routing strategies generally remained for the peripheral nodes (preferred NAV), satellite connectors (preferred DIF), connector hubs (preferred SP), and provincial hubs (only used SP and NAV), with the strength of preference varied slightly (Fig. S4D). Taken together, we could conclude that our main findings were robust and reproducible.

## 3. Discussion

This study proposed a communication model for the human brain structural connectivity network (SCN) based on the idea of region-wise hybrid routing, which allowed brain regions to adopt different routing strategies for neural signal transmission. The hybrid routing strategy was approximated through a genetic algorithm (GA) that aimed at maximizing the coupling between structural routing efficiency and functional connectivity. Based on the obtained solution (namely ‘HYB’), our main findings were as follows. Firstly, the regions adopting different strategies did not randomly scatter over the brain but showed a specific clustering pattern that partially complied with the partition of functional modules. Secondly, the proposed HYB strategy was advantageous over the other three strategies both in terms of capturing the FCN and supporting robust SCN communication. Thirdly, brain regions in lower-order functional modules (e.g., VIS and SMN) inclined to choose the routing strategies requiring more global information (global topology for SP and spatial embedding for NAV), while those in higher-order functional components increased the usage of DIF. In addition, compared to the brain regions using SP and NAV, the brain regions using DIF had more structural links, participated in more functional modules, but played a less dominant role within modules. In conclusion, our results supported that the preference for global information utilization had regional heterogeneity. The resulting communication model facilitated the integration and segregation of information and thus the formation of functional modules.

## Supporting information

routing_fig

---

**Figure S2.**
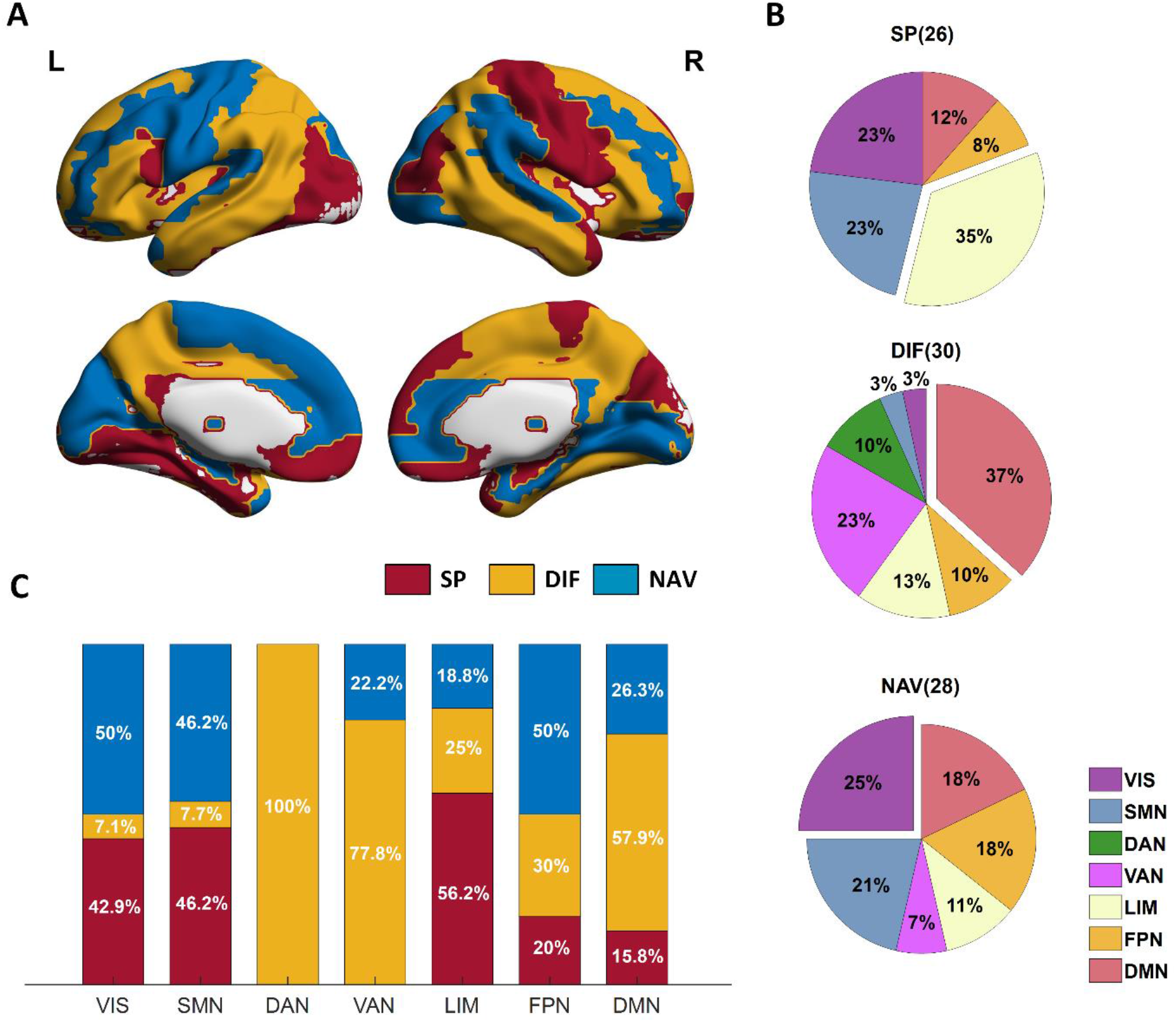
Hybrid routing strategy (B-HYB) found on the validation dataset BNU. A: Brain map of B-HYB. B: Modular distribution of regions using each routing strategy, where the number in the parentheses indicates the number of regions choosing the strategy. C: Proportions of the three routing strategies used by regions in each functional module. The functional modules were derived from Yeo *et al*. [29].

**Figure S3.**
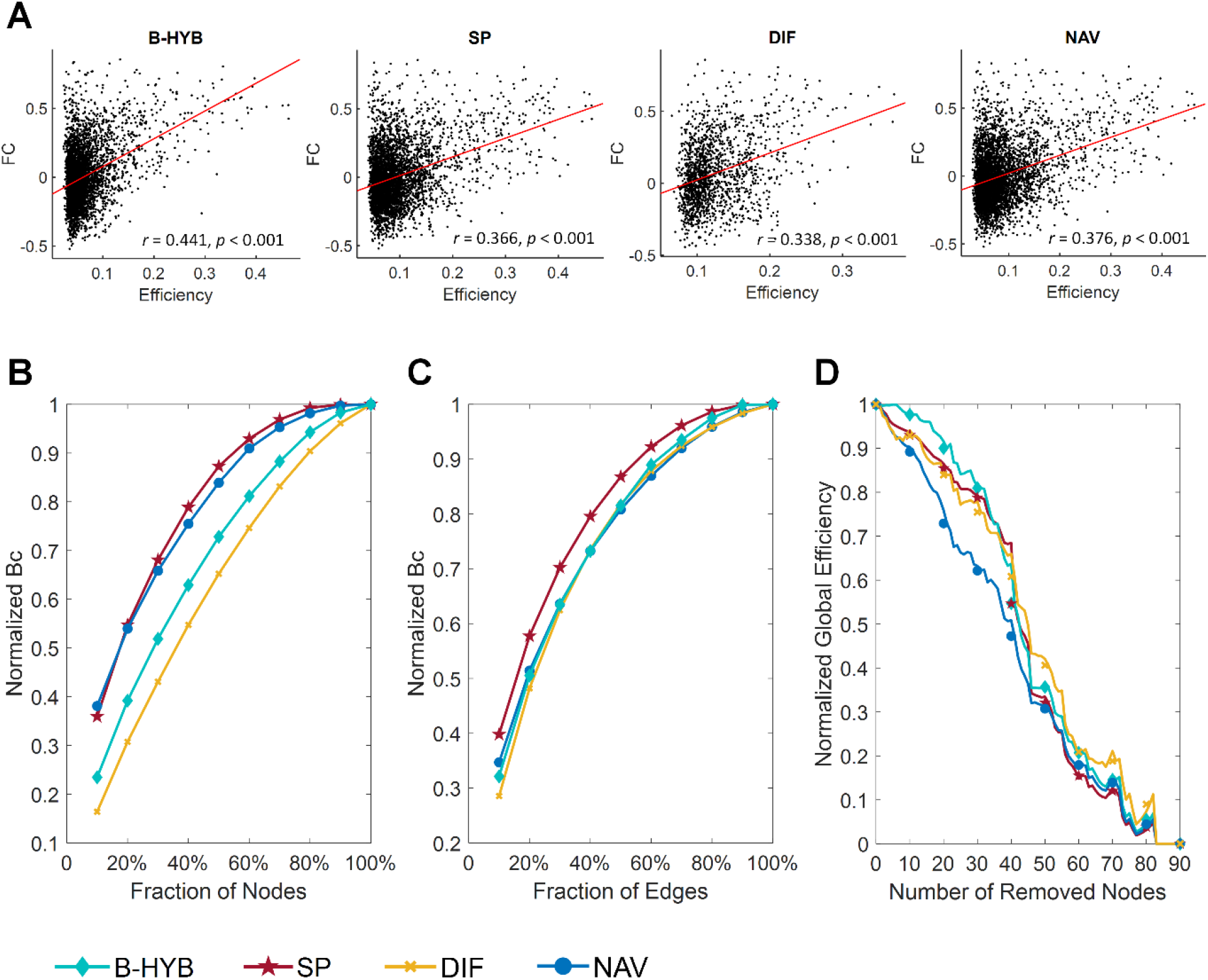
Comparison of the proposed hybrid routing strategy (B-HYB) and the three typical classic routing strategies (SP, DIF, and NAV) on the BNU dataset. A: Comparison of the global-level correlation between FC and routing efficiency. B: Comparison of normalized nodal betweenness; C: Comparison of normalized edgewise betweenness; D: Comparison of relative global network efficiency during targeted attack.

**Figure S4.**
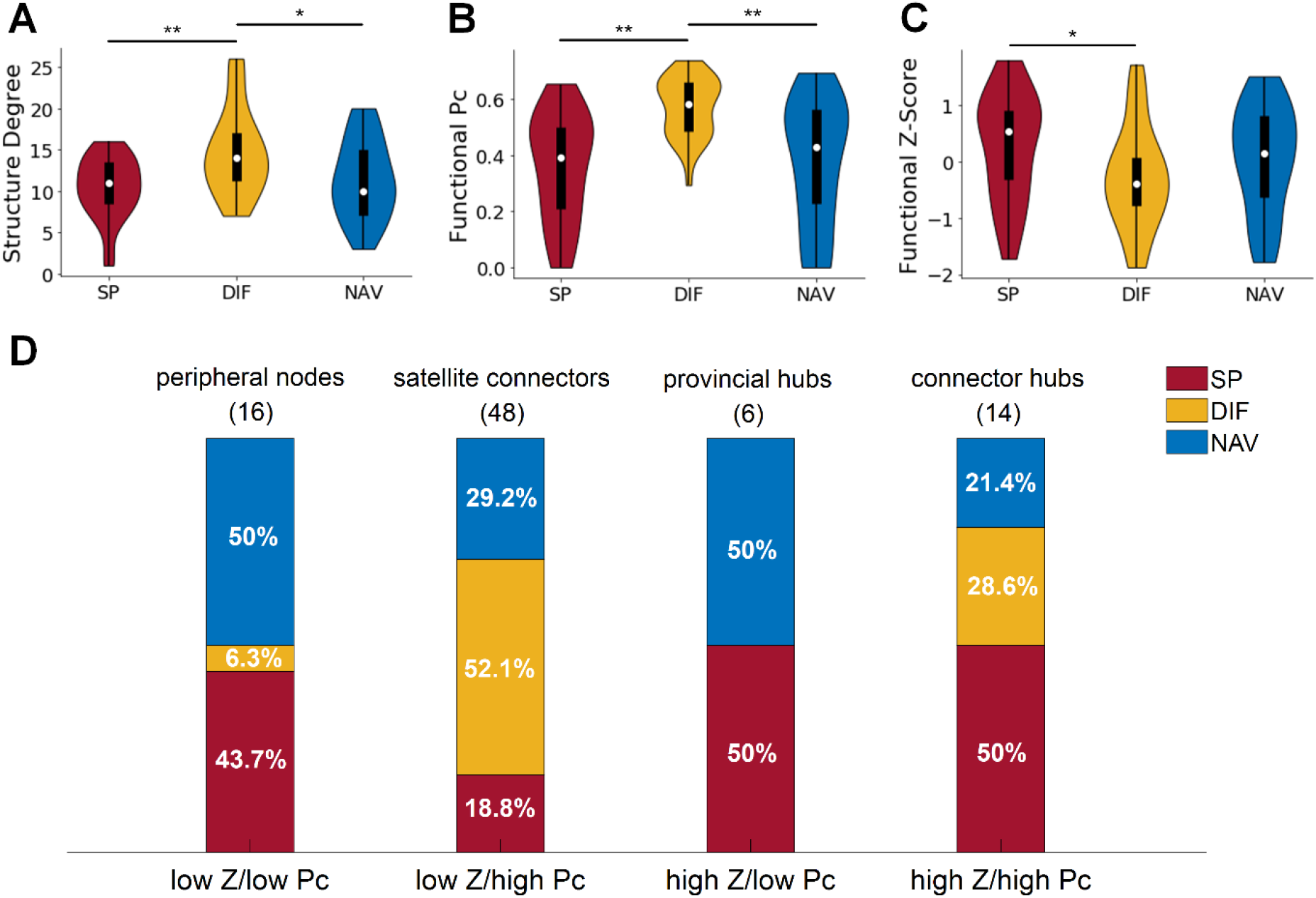
Plausibility analysis on the regional choices of routing strategies in B-HYB. A: Comparison of structural degree. B: Comparison of functional participation coefficient (Pc). C: Comparison of functional within-module degree Z-score (Z). D: Distributions of routing strategies adopted by the four types of regions classified according to the Pc-Z coordinate, with Pc > 0.3 corresponding to ‘high Pc’ and Z > 0.9 corresponding to ‘high Z’.

