## Supplementary figures and images for "A Hybrid Communication Pattern in Human Brain Structural Network Revealed by Evolutionary Computation"

### routing_fig

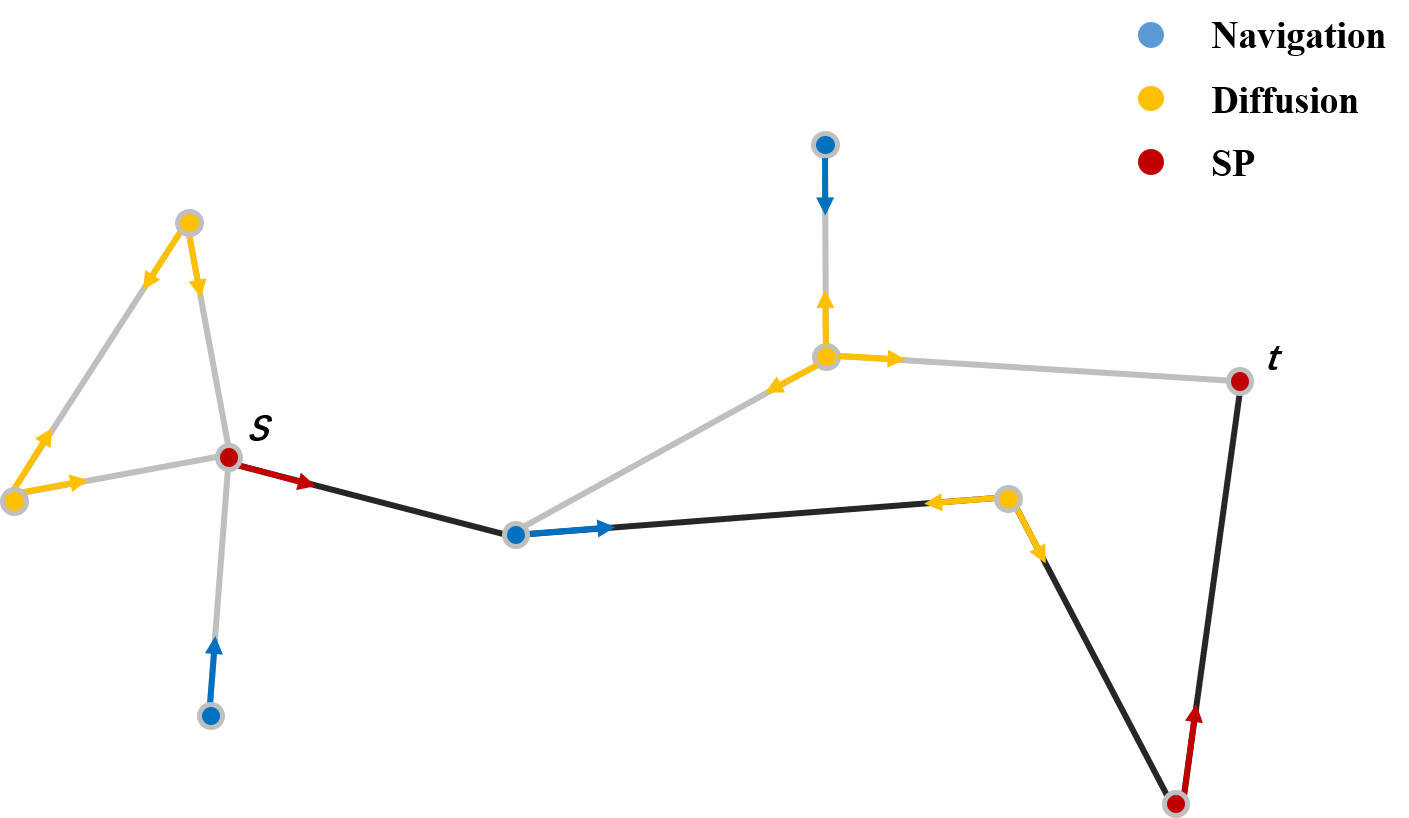
